# Non-Physiological Potassium Concentrations in Commercial Culture Media Trigger Acute Epileptiform Activity in Human iPSC-Derived Neurons

**DOI:** 10.1101/2025.05.31.657142

**Authors:** Tim Lyckenvik, Julia Izsak, Erik Arthursson, My Forsberg, Kalle Johansson, Henrik Zetterberg, Markus Axelsson, Pontus Wasling, Eric Hanse, Stephan Theiss, Sebastian Illes

## Abstract

Neuronal *in vitro* cultures are pivotal for studying brain electrophysiological function and dysfunction. Neuronal activity and communication are regulated by extracellular ion concentrations. Therefore, cell culture medium ion concentrations should ideally mimic those of cerebrospinal fluid (CSF) – considered as the milieu for brain cells *in vivo*.

In this study, we demonstrate that commonly used cell culture media, including Neurobasal™ (+/- A), Neurobasal Plus™, and BrainPhys™ media, do not accurately replicate human CSF ion concentrations. Using human iPSC-derived neuronal networks on microelectrode arrays, we show that the abnormally high potassium concentrations present in all tested cell culture media induce acute epileptiform activity, similar to that elicited by the convulsive drug 4- aminopyridine.

These findings raise a critical question: How can human *in vitro* neuronal activity be defined as physiological and reliably distinguished from pathophysiological activity, if the routinely used ion concentrations in *in vitro* experiments are causing aberrant neuronal activity?

**Summary:** The neuronal activity in neuronal *in vitro* culture relies on extracellular ion concentrations, which should mimic cerebrospinal fluid (CSF). This study shows that common cell culture media and widely used artificial CSF composition in neuroscience research fail to replicate CSF ion levels, causing non-physiological and rather pathological neuronal activity.

## Introduction

Electrophysiology methods applied to *in vitro* brain cell cultures have substantially contributed to understanding the principles of brain function and dysfunction. Like neurons in the brain, *in vitro* neurons are electrically active, communicate via chemical and electrical synapses, and self-organize into neuronal networks generating spontaneous synchronized activity and local field potentials, enabling the study of neuronal function at both the micro- and mesoscale.

Both passive and active electrophysiological properties of a neuron are determined by its extracellular ion concentrations—primarily sodium, chloride, potassium, magnesium, and calcium ions. Exposing neurons to a simple buffer solution containing only these ions, along with glucose and bicarbonate, in *in vitro* brain slice and neuronal culture experiments allows for the study of passive and active electrophysiological properties of neurons. Thus, this approach is the standard *in vitro* method in neuroscience. Because the ion concentrations in this buffer solution are based on known cerebrospinal fluid (CSF) levels, it is referred to as artificial cerebrospinal fluid (aCSF). Since aCSF alone cannot support long-term survival of neurons *in vitro*, specialized cell culture media are used for *in vitro* neural cultures. These cell culture media contain the same core ions as aCSF and are supplemented with vitamins, amino acids, lipids, antioxidants and other ions. Ideally, these media should replicate the ion composition of CSF to maintain physiological relevance.

While researchers typically prepare their own aCSF in the lab, neuronal cell culture media are exclusively purchased as ready-to-use solutions. Since the introduction of Neurobasal™ as the first chemically defined cell culture medium that allowed *in vitro* neurons to survive without fetal serum in the 1990s^1^, Neurobasal™ (licenced by Thermo Fisher) and BrainPhys™ (licenced by StemCell Technologies) have become the most widely used media in neuroscience research today. Originally, Neurobasal™ and DMEM/F12 media—often supplemented with serum—were used to culture rodent and human-induced pluripotent stem cell (hiPSC)-derived neurons. However, in a seminal study by Bardy et al. (2015), these media were found to be inferior to an aCSF whose ion concentrations were intended to more closely match those of human CSF in supporting neuronal electrophysiological activity at synaptic, single-cell, and network levels^2^.

Building on these findings, Bardy et al. developed a new cell culture medium, BrainPhys™, by adjusting sodium and calcium levels, removing certain neuroactive amino acids, and modifying glucose and osmolarity levels^2^. These modifications improved the electrophysiological functionality of rodent and human iPSC-derived neurons, establishing BrainPhys™ as a widely used cell culture medium in neuroscience and stem cell research. More recently, Neurobasal™ Plus has been introduced, to enhance the original Neurobasal™ medium. However, its exact composition remains proprietary. Neurons cultured in Neurobasal™ Plus reportedly exhibit accelerated neurite growth, synapse development, improved cell health, and increased neuronal network activity ^3^.

The importance of having brain specific ion concentrations becomes evident when considering that CSF and serum have different concentrations of potassium, magnesium, sodium, calcium, and chloride. Previously, we performed a person-specific analysis of CSF-serum ion concentration correlations by measuring ion levels in cerebrospinal fluid and blood serum samples collected from the same individuals^4^. Because CSF freely communicates with brain interstitial fluid (ISF)^5, 6^, it serves as a valuable proxy for assessing the composition of the brain’s extracellular environment. Our analysis revealed no correlation between potassium, chloride, or magnesium concentrations in CSF and serum, and a moderate correlation for sodium and calcium^4^. Specifically, our findings, along with previous studies^7–18^, suggest the presence of regulatory mechanisms within the brain or at the blood-brain barrier that lower serum potassium concentrations from approximately 4.1 mM to about 2.9 mM in CSF, while conversely increasing serum magnesium concentrations from around 0.9 mM to approximately 1.14 mM in CSF^4^.

Given that elevated potassium levels increase neuronal excitability, and that reduced magnesium levels enhance NMDA receptor activity—among other effects—these regulatory mechanisms suggest that the brain actively creates a specific ionic environment in the CSF to keep neuronal activity at a certain level and prevent excessive neuronal activity that would otherwise occur if neurons were directly exposed to serum ion concentrations.

Considering that the brain precisely regulates CSF ion concentrations—and that sodium, chloride, potassium, magnesium, and calcium are critical for defining the passive and active electrophysiological properties of neurons—we investigated, in the present study, whether commonly used culture media (BrainPhys™, Neurobasal™ Plus, and DMEM/F12) accurately reflect physiological CSF ion concentrations. If discrepancies were found, we aimed to examine their impact on in vitro neuronal function using human iPSC-derived neurons and microelectrode array (MEA) technology—a state-of-the-art platform for studying human neuronal electrophysiology at the mesoscale.

## STAR Methods

### Ethical statement

For CSF sampling, all subjects participated in the study voluntarily and provided written consent. CSF sampling was approved by the Swedish Ethical Review Authority in Gothenburg (#223-15, #492-18, #942-12).

### Cultivation, neural differentiation of human iPSC lines and 3D neural aggregate formation

The iPSC line (ChIPSC4, Takara Bio Europe AB (formerly Cellartis AB)) was cultured under feeder-free conditions in Cellartis DEF-CS™ (Takara Bio Europe AB) or mTESR1 (StemCellTechnologies) at 37°C in a humidified atmosphere of 5% CO2 in air. By applying the DUAL-SMAD inhibition protocol^19^ frozen stocks of human iPSC-neural stem cells (hiPSC-NSC) were obtained according our previously published procedure ^20^. Frozen stocks of hiPSC-NSC were thawed and cultured in neural culture media on Poly-L-Ornithine (PLO) /Laminin-coated 3.5 cm culture plates. Neural culture media consisted of DMEM/F12 GlutaMAX, Neurobasal™, N2 supplement, B27 supplement, 5µg ml-1 insulin, 1 mM Ultra glutamine, 100 µM non-essential amino acids, 100 µM 2-mercaptoethanol, 50 U ml-1 penicillin and streptomycin. After 7 to 10 days, four to eight hiPSC-3D-neural aggregates were seeded as a 5 µl drop directly on PLO/laminin-coated multi-electrode arrays (MEAs). For neuronal differentiation, BrainPhys™-media supplemented with N2 supplement, B27 with vitamin A, 2 mM Ultra glutamine, 50 U ml^-1^ Pen/Strep, and 200 µM ascorbic acid were used. The media was further supplemented with neurotrophic factors: brain derived neurotrophic factor (BDNF), glial derived neurotrophic factor (GDNF), transforming growth factor-β (TGF-β) [20 ng/ml each], and optional DAPT [10 µM]. Half media exchanges were performed twice a week. For further information about used culture media and procedures see elsewhere^20^.

### Multi-electrode array recordings and experiments

MEAs had Ti/TiAu electrodes with PEDOT-CNT (carbon nanotube poly-3,4-ethylene- dioxythiophene) of 30 µm diameter and 200 µm spacing. Electrode configuration was 9 recording electrodes per well (6-well MEAs) or 3 recording electrodes per well (96-well MEAs). MEA-electrodes had an input impedance of 30–50 kΩ according to the specifications of the manufacturer (Multi Channel Systems). The recording sampling rate was 25 kHz on all MEA electrodes using a MEA2100 system (Multi Channel Systems). MC_Rack (Multi Channel Systems) and Multi Channel Experimenter (version 2.20.0.22133; Multi Channel Systems) were used to visualize and store MEA data. Offline-spike detection and synchronous network activity were performed by the SPANNER software suite (RESULT Medical) and Multi Channel Analyzer (version 2.20.2.22291; Multi Channel Systems).

Raw data was filtered using a Butterworth high pass filter of the 2^nd^ order with cutoff at 200Hz. Spike detection was done by the software using a threshold of 4-5 standard deviations of the noise of each channel respectively, for both the rising and falling edge, during a non-network bursting period with low spike rate in the medium with the least amount of activity, and kept at this absolute threshold throughout all conditions for each culture respectively. Dead time was set to 3ms, (1ms pre, and 2ms post trigger).

The following network burst detection parameters were used (in ms): Max. interval to start burst: 50; Max. interval to end burst: 50; Min. interval between bursts: 100-500; Min. duration of burst: 50; Min. number of spikes in bursts: 5-50. In ten BrainPhys™ recordings, an increase in noise levels was observed following BrainPhys™ application. To minimize false-positive burst detection, the minimum spike threshold for network burst detection was accordingly increased for these recordings.

Human iPSC-derived neural cultures were maintained in BrainPhys™ medium with standard supplements until the day of the experiment. For the comparison of 2.8, and 4 mM K^+^, the cultivation media was replaced with 100 µl aCSF containing ion concentrations matching those of hCSF, and MEA recordings were conducted. Afterward, a potassium chloride solution was added to achieve an aCSF with a final potassium concentration of 4 mM, followed by additional MEA recordings.

For the other experiments, the cultivation media was removed, and either aCSF with varying potassium concentrations (as detailed in the results), human CSF, or fresh BrainPhys™ medium (100 µl per well of a 6-well MEA) was added.

MEA recordings were performed for 6 minutes under all experimental conditions. For quantitative analyses, the last two minutes of the recordings of the experiments comparing aCSF with different levels of K^+^, while all 6 minutes of human CSF experiments were analysed.

The ion-matched aCSF was prepared to match the previously determined concentrations of the major ions (sodium, potassium, magnesium, calcium and chloride), bicarbonate and glucose in human CSF^21^. The contents of the ion-matched aCSF were (in mM): 124.2 NaCl, 2.79 KCl, 1.14 MgCl, 23 NaHCO, 0.4 NaH 2 PO4, 1.18 CaCl, and 3.66 D-glucose.

For the 4-AP experiments, epileptiform activity was induced by adding 4-AP to the cell culture media at a final concentration of 100 µM.

### CSF collection and analysis

28 healthy volunteers were recruited from the local community for two previous studies^22, 23^. Prior to inclusion, all subjects underwent screening for the exclusion criteria of symptoms or diagnosis potentially related to the CNS, intake of medications or drugs, and abnormal sleep habits or general health. All CSF samples were obtained by routine lumbar puncture performed by an experienced neurologist using an atraumatic needle (22-25G). The CSF samples were centrifuged at 2000 g for 10 minutes at room temperature to remove cells and debris and supernatant were stored in aliquots of 1ml at −80°C pending biochemical analysis and experimental use.

The measurements were carried out by board-certified laboratory technicians at the Clinical Chemistry Laboratory at Sahlgrenska University Hospital using accredited methods with inter-assay coefficients of variation below 2%. The laboratory is accredited under the Swedish Accreditation body (Swedac).

Concentrations of Na^+^, K^+^ and Cl^-^ were measured using ion-selective electrodes (ISEs), integrated into the Cobas c 501 instrument, which have been approved for clinical use without clinically relevant interferences (Roche Diagnostics). ISE Standards Low (S1) and High (S2) are used to calibrate the methods at least once per day (Roche Diagnostics). Ca^2+^ and Mg^2+^ concentrations were measured by colorimetric o-cresolphthaleion and chlorophonazo III methods, respectively, in the Cobas c 501 instrument, according to instructions from the manufacturer (Roche Diagnostics). The methods are calibrated after each reagent lot change, using Standards Low (S1) and High (S2) for Ca^2+^ and Mg^2+^, respectively (Roche Diagnostics).

### Statistical analysis

For statistical analysis GraphPad Prism (version 10.4.0) was used. Wilcoxon matched pairs signed rank test was used to calculate the shown *p* values. The data are presented as mean values ± standard deviation (SD). *n* refers to the number of technical replicates, and *p*-values below 0.05 are considered significant. All this information is provided within the figures. For detailed values per experiment, see Supplementary Table 2.

### Resource Availability

All data reported in this paper will be shared by the lead contact upon reasonable request.

## Results

### Lack of standardization in aCSF ion composition across neuroelectrophysiological studies

Prior to conducting experimental work, we reviewed the literature to gain insights into the typical ion concentrations used in *in vitro* neuroelectrophysiology experiments, including primary neuronal cultures and both animal and human brain slice preparations.

Typical aCSF is composed of sodium (Na⁺), chloride (Cl⁻), potassium (K⁺), calcium (Ca²⁺), and magnesium (Mg²⁺) ions, as these are the major ions regulating neuronal excitability and synaptic communication. Supplementary Table 1 provides numerous examples of research articles where aCSF have been prepared and used for neuro-electrophysiology experiments, highlighting a common phenomenon—high potassium concentrations (>3 mM) in aCSF used for acute neuroelectrophysiology recordings.

While this is not a comprehensive assessment of the field, we consider it sufficient to demonstrate the lack of consensus within neuroscience research regarding appropriate ion concentrations, as the composition of aCSF used in *in vitro* and *ex vivo* experiments varies widely.

### Commercial cell culture media fails to replicate human CSF ion composition, with markedly elevated potassium levels

Given the wide range of ion concentration compositions used in aCSF for acute neurophysiological experiments (Suppl. Table 1), we measured and compared the concentrations of Na^+^, Cl^-^, K^+^, Mg^2+^ and Ca^2+^ in human CSF samples obtained from 28 young and healthy volunteers (mean age 25.3 ± 5.3) with measured ion concentrations in BrainPhys™, Neurobasal PLUS™, and DMEM/F12. Our analysis revealed that none of the commercial cell culture media matched the ionic composition of hCSF (Table 1). In particular, the potassium concentrations were substantially higher, while magnesium levels were lower, across all three media. BrainPhys™ and DMEM/F12 exhibited sodium and chloride levels similar to hCSF, but calcium concentrations were lower. Neurobasal™ displayed pronounced deviations from hCSF across all measured ions.

**Table 1.**
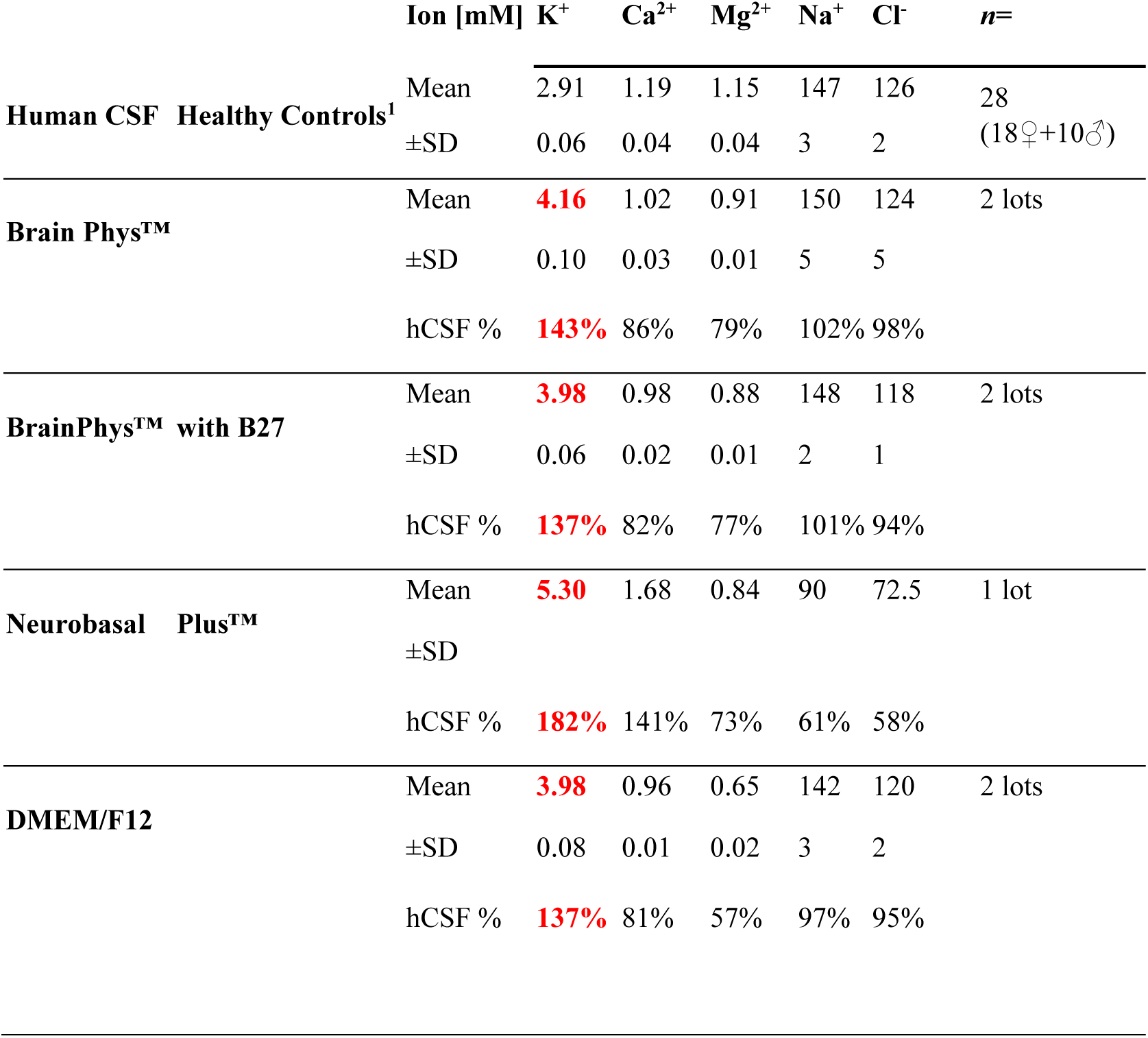
Table summarizes the concentrations of potassium, calcium, magnesium, sodium, and chloride ions in human cerebrospinal fluid samples, and commercially available neural cell culture media and BrainPhys-based and Neurobasal Plus-based condition media (collected after 3 days of a full media exchange). Abbreviations: SD, standard deviation;hCSF %, proportion of the typical human cerebrospinal fluid concentration.

|  |  | Ion [mM] | K<sup>+</sup> | Ca<sup>2+</sup> | Mg<sup>2+</sup> | Na<sup>+</sup> | Cl<sup>-</sup> | n= |
| --- | --- | --- | --- | --- | --- | --- | --- | --- |
| Human CSF | Healthy Controls<sup>1</sup> | Mean | 2.91 | 1.19 | 1.15 | 147 | 126 | 28 |
|  |  | ±SD | 0.06 | 0.04 | 0.04 | 3 | 2 | (18♀+10♂) |
| Brain Phys™ |  | Mean | 4.16 | 1.02 | 0.91 | 150 | 124 | 2 lots |
|  |  | ±SD | 0.10 | 0.03 | 0.01 | 5 | 5 |  |
|  |  | hCSF % | 143% | 86% | 79% | 102% | 98% |  |
| BrainPhys™ with B27 |  | Mean | 3.98 | 0.98 | 0.88 | 148 | 118 | 2 lots |
|  |  | ±SD | 0.06 | 0.02 | 0.01 | 2 | 1 |  |
|  |  | hCSF % | 137% | 82% | 77% | 101% | 94% |  |
| Neurobasal Plus™ |  | Mean | 5.30 | 1.68 | 0.84 | 90 | 72.5 | 1 lot |
|  |  | ±SD |  |  |  |  |  |  |
|  |  | hCSF % | 182% | 141% | 73% | 61% | 58% |  |
| DMEM/F12 |  | Mean | 3.98 | 0.96 | 0.65 | 142 | 120 | 2 lots |
|  |  | ±SD | 0.08 | 0.01 | 0.02 | 3 | 2 |  |
|  |  | hCSF % | 137% | 81% | 57% | 97% | 95% |  |

### Unphysiological artificial cerebrospinal fluid with 4 mM potassium concentration causes acute epileptiform activity in human neurons

Given the discrepancy between potassium levels in hCSF and commonly used cell culture media and aCSF in *in vitro* electrophysiology experiments, we assessed if the neuronal network activity differed between human neurons exposed to aCSF with physiological (2.9 mM) or elevated (4 mM) potassium concentration. We used human iPSC neurons with partial or synchronous neuronal network activity—characterized by simultaneous population bursts across different electrodes— and exposed them sequentially to aCSF with 2.9 mM and 4 mM K⁺ concentration.

We observed that elevation to 4 mM K⁺ caused an immediate and prominent increase in neuronal network activity. Within seconds, the networks exhibited highly synchronous, rhythmic activity, characterized by a significant increase in network burst rate and a shortening of burst duration, while the number of spikes per minute concomitantly increased (Figure 1A, right panel, gray traces). Over the course of the four-minute adaption period, the network burst duration gradually lengthened while the network burst rate decreased; however, network burst rate stabilized and remained substantially higher than under 2.9 mM K⁺.

**Figure 1.**
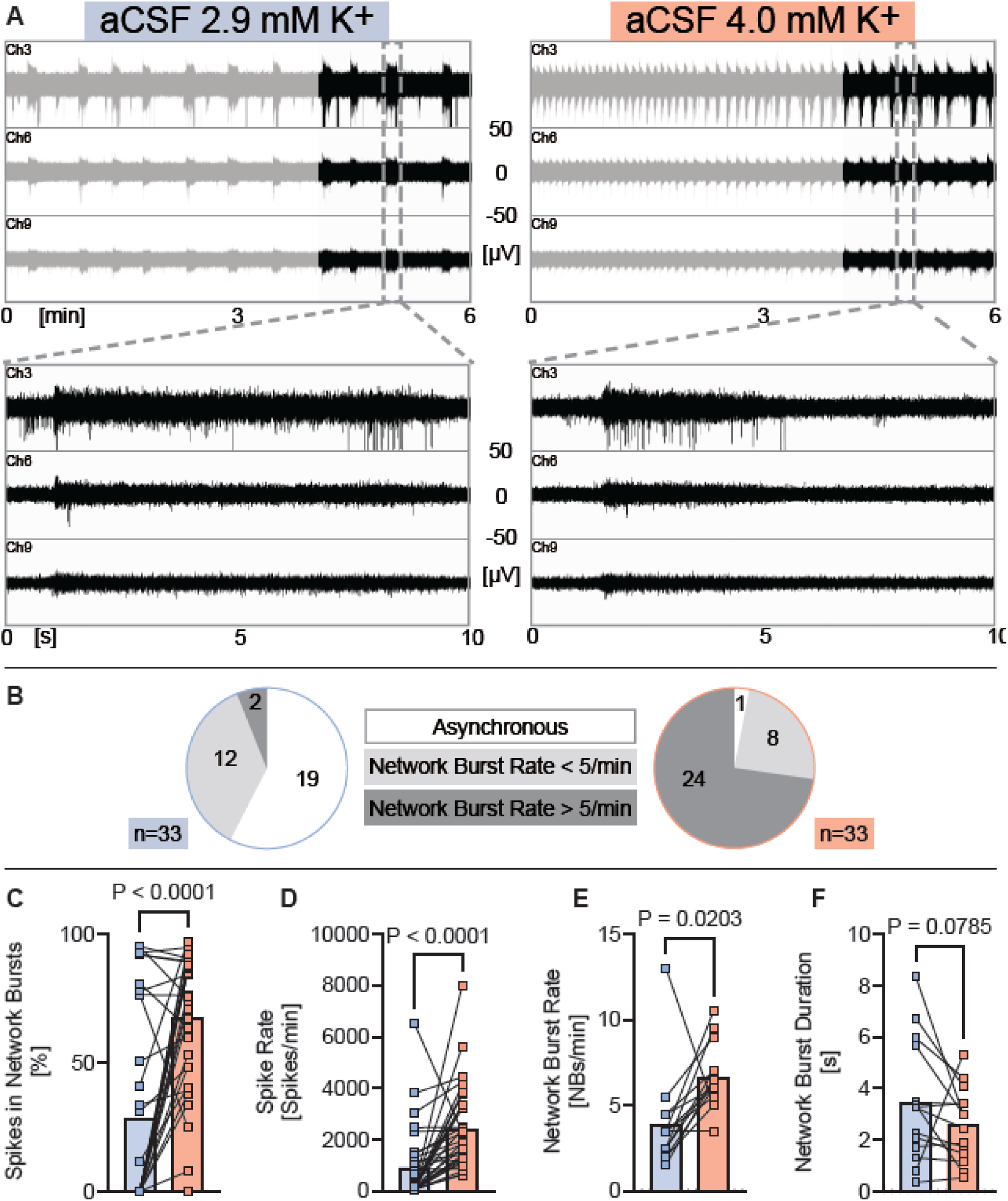
Unphysiological aCSF with 4 mM potassium concentrations causes epileptiform activity in human neurons. (**A**) Representative examples of MEA-recordings (three individual channels are shown) showing neuronal network activity with a time resolution of 6 minutes and 10 seconds of human iPSC neurons exposed to aCSF with either 2.9 mM or 4 mM potassium. (**B**) Diagrams showing the proportion of experiments where network burst rate was 0 (= asynchronous), under five per minute or over five per minute, recorded from human iPSC neurons exposed to aCSF with either 2.9 mM or 4 mM potassium. (**C-F**) Diagrams illustrating the change of neuronal network parameters under each condition respectively. Individual mean values and *p* values from Wilcoxon tests are shown. Experiments were conducted between 48 to 77 days *in vitro*. Mean, SD and *n* are presented in Supplementary Table 2.

To capture stable activity patterns, we quantified data from the final two minutes of each recording. During this period, 18 of the 19 networks that lacked synchronous network bursts in 2.9 mM K^+^, had transitioned to exhibiting synchronous bursting activity in 4.0 mM K^+^ (Figure 1B).

Comparing the neuronal network properties during the last two minutes under 2.9 mM and 4 mM potassium showed that 4 mM potassium substantially increased spike synchrony (% of spikes organized into network bursts) and spike rate (Figure 1C, D). Among the 14 cultures that exhibited network bursts under both 2.9 mM and 4 mM K⁺, we observed that 4 mM K⁺ elicited a strong increase in network burst rate, with a nonsignificant tendency toward shorter burst duration (Figure 1E, F). The activity pattern elicited by 4 mM potassium resembled epileptiform activity, prompting us to conduct experiments using 4-aminopyridine (4-AP), a non-selective voltage-dependent K⁺ channel blocker commonly used to induce epileptiform activity in acute brain slice preparations^24^, rodent^25^ and human iPSC neuronal^26^ *in vitro* models. Indeed, 4-AP elicited highly synchronous, rhythmic and repetitive bursts of activity with high burst rate (Suppl. Figure 1), closely resembling the activity induced by 4mM potassium. Thus, elevating K^+^ concentration by just 1.1mM in aCSF, from 2.9 to 4.0 mM, is sufficient to induce activity patterns resembling drug-induced epileptiform activity in human neuronal networks.

### Human neurons exposed to BrainPhys™ show epileptiform activity similar to artificial cerebrospinal fluid with 4 mM potassium

Since our ion concentration measurements of BrainPhys™ samples revealed an elevated potassium concentration of 4 mM, we compared the neuronal network activity of human iPSC neurons consecutively exposed to aCSF with 4 mM potassium to the activity after applying BrainPhys™ to the same cultures (Figure 2). Interestingly, the application of BrainPhys™ led directly to a neuronal network activity level similar to that elicited by 4 mM potassium in aCSF. No dynamic changes in network bursting pattern were observed during the six-minute BrainPhys™ recordings. However, for consistency, we analyzed the last two minutes of recordings under both conditions. Across both groups, only one culture under BrainPhys™ showed an absence of network bursts, and the network burst rate was generally above five network bursts per minute under both conditions (Figure 2B).

**Figure 2.**
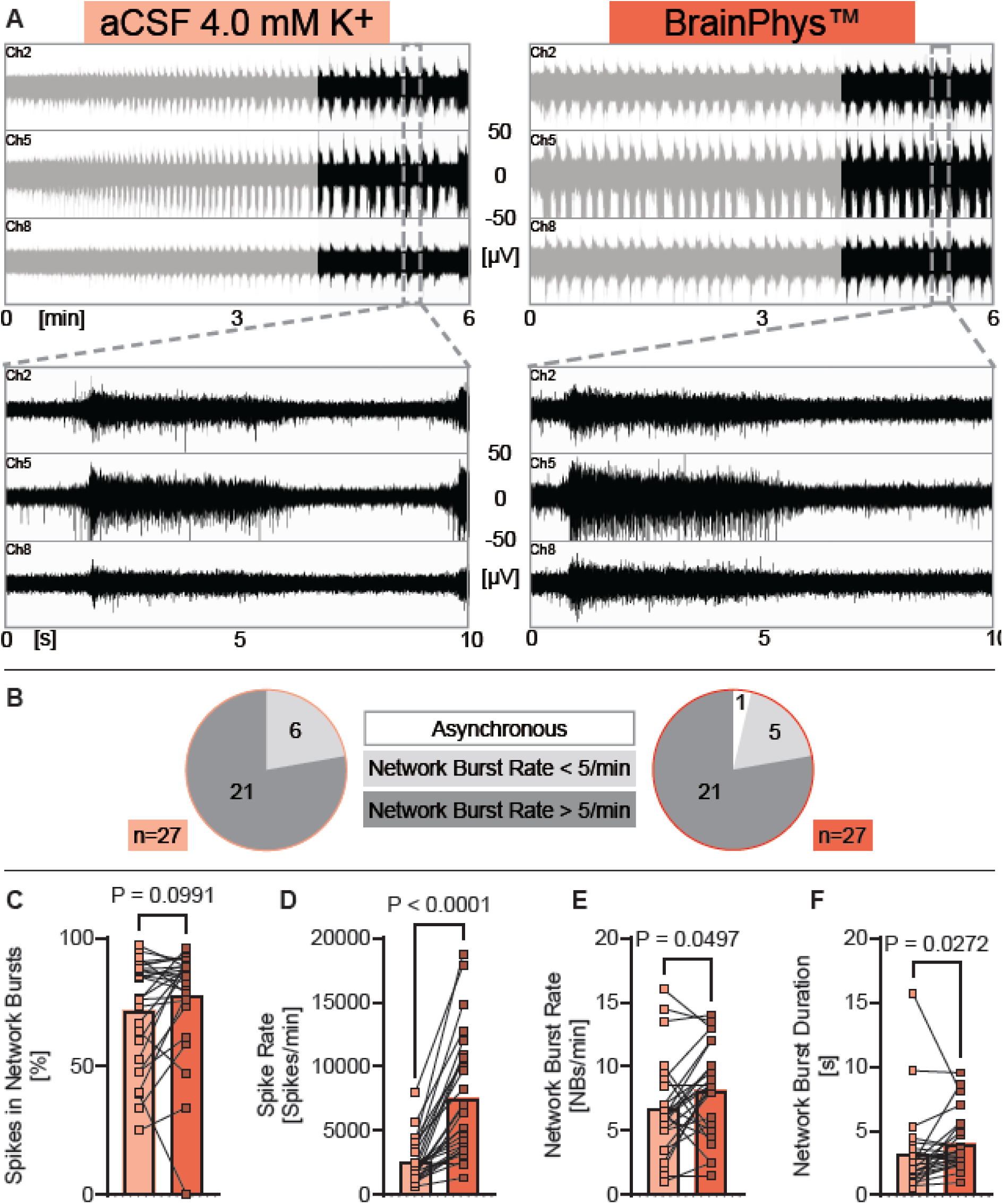
Human neurons exposed to BrainPhys show epileptiform activity similar to that elicited by aCSF with 4 mM potassium. (**A**) Representative examples of MEA-recordings (three individual channels are shown) showing neuronal network activity of human iPSC neurons exposed either to aCSF with 4 mM potassium or to BrainPhys™ with a time resolution of 6 minutes, and 10 seconds of human iPSC neurons exposed either to aCSF with 4 mM potassium or to BrainPhys™. (**B**) Circle diagrams showing the proportion of experiments where network burst rate was 0 (= asynchronous), under five per minute, or over five per minute, recorded from human iPSC neurons exposed either to aCSF with 4 mM potassium or to BrainPhys™. (**C-F**) Diagrams illustrating the change of neuronal network parameters under each condition respectively. Individual mean values and *p* values from Wilcoxon tests are shown. Experiments were conducted between 48 to 77 days *in vitro*. Mean, SD and *n* are presented in Supplementary Table 2.

The network burst rate was slightly higher (Figure 2E), and the network burst duration was slightly longer in BrainPhys™ (Figure 2F). Spike synchrony was similar in both conditions (Figure 2C), while the spike rate was substantially higher in BrainPhys™ (Figure 2D).

These findings indicate that BrainPhys™ induces longer network bursts and increases spiking activity, further amplifying neuronal activity compared to the elevated activity elicited by supraphysiological K^+^ levels alone. Rather than mimicking the K^+^-induced epileptiform activity, BrainPhys™ appears to exacerbate it.

### Acute application of human CSF increases neuronal network activity in comparison to physiological aCSF

Next, we compared the neuronal activity of human *in vitro* neurons exposed to human CSF versus exposure to aCSF with identical ion concentrations of Ca^2+^, Mg^2+^, Na^+^, K^+^ and Cl^-^. While we have shown that human iPSC neurons cultured in human CSF accelerated several maturation processes and exhibited increased neuronal network activity^27^, the acute and direct effects of human CSF, compared to artificial CSF with ion concentrations matched to hCSF, on human iPSC-derived neuronal networks had not been previously assessed.

To address this, we exposed human iPSC-derived neuronal networks to physiological aCSF, then switched to human CSF, and then switched back to aCSF. After removing hCSF, a three-time wash with aCSF was performed.

Human iPSC-derived neurons exposed to hCSF displayed increased spike synchrony (measured as spikes organized into network bursts), spike rate and network burst rate, compared to aCSF (Figure 3C-F). However, the burst duration was not significantly different (Figure 3F). Additionally, 75% of the 40 analysed cultures exhibited network bursts under hCSF, whereas only 43% showed network bursts under physiological aCSF (Figure 3B).

**Figure 3.**
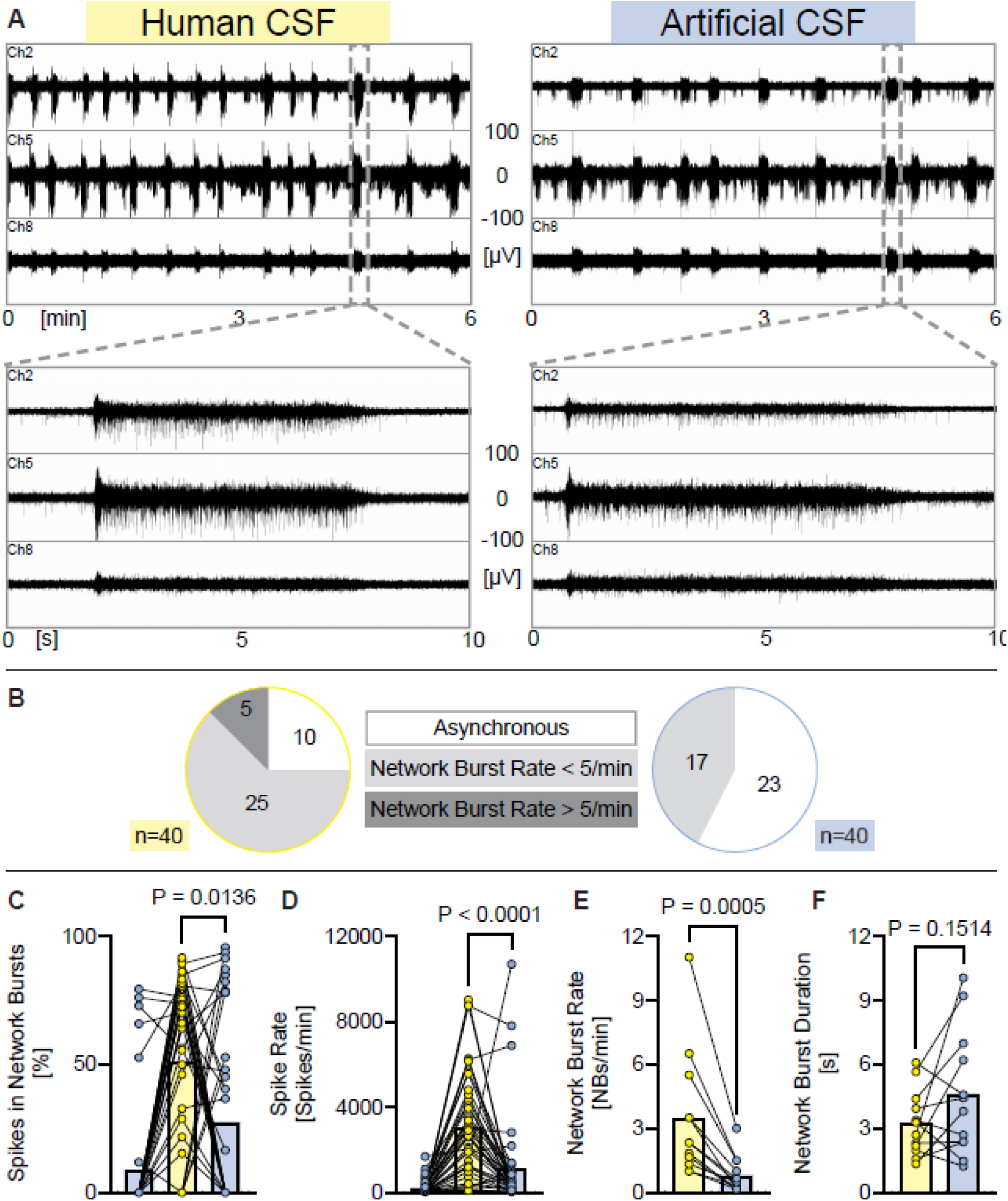
Acute application of human CSF increases neuronal network activity in comparison to aCSF with ion concentrations matched to human CSF. (**A**) Representative examples of MEA-recordings (three individual channels are shown) showing neuronal network activity with a time resolution of 6 minutes, and 10 seconds of human iPSC neurons exposed either to human CSF or to artificial CSF with ion concentrations matched to typical hCSF ion concentrations. (**B**) Diagrams showing the proportion of experiments where network burst rate was 0 (= asynchronous), under five per minute, or over five per minute, recorded from human iPSC neurons exposed either to hCSF or to aCSF. (**C-F**) Diagrams illustrating the change of neuronal network parameters under each condition respectively. Individual mean values and *p* values from Wilcoxon tests are shown. Experiments were conducted between 48 to 77 days *in vitro*. Mean, SD and *n* are presented in Supplementary Table 2. Note: each data point in the hCSF data set represents the activity of an individual human iPSC-neuronal culture exposed to a hCSF sample obtained from a human individual.

In summary, human iPSC neurons acutely exposed to hCSF exhibit higher neuronal network activity compared to those exposed to aCSF with ion concentrations, glucose, and bicarbonate levels matched to hCSF.

### BrainPhys™ increases spontaneous activity in human iPSC-derived neuronal networks compared to human cerebrospinal fluid

Since both hCSF and BrainPhys elicit higher activity compared to aCSF with physiological ion concentrations, making a direct conclusion difficult, we conducted an experiment comparing the neuronal network activity of the same human iPSC neurons first exposed to human CSF and subsequently exposed to BrainPhys™ medium, with aCSF washing steps in between (Figure 4A).

**Figure 4.**
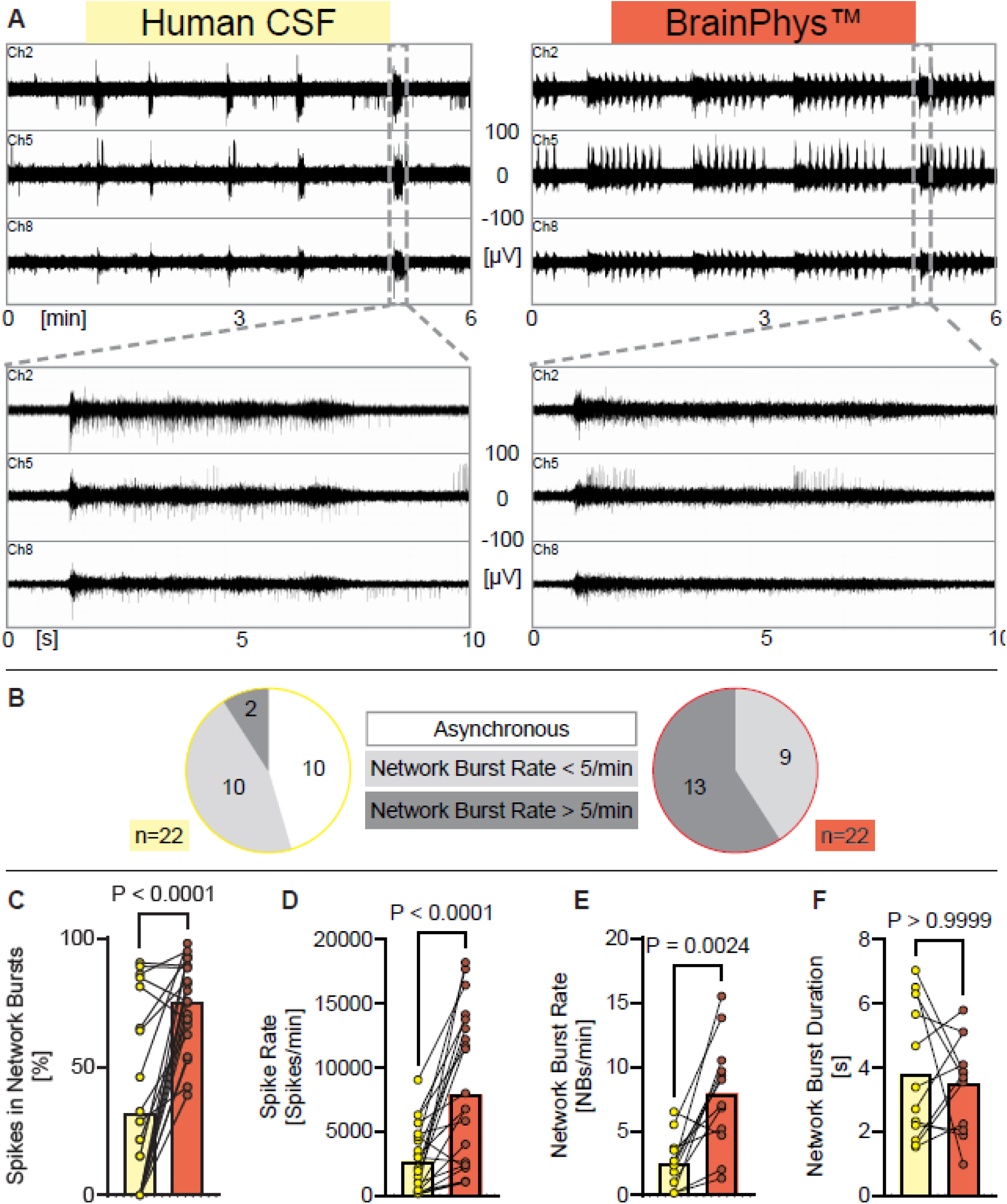
Human iPSC neurons exposed to human CSF show different synchronous neuronal network activity as elicit by BrainPhys™. (**A**) Representative examples of MEA-recordings (three individual channels are shown) showing neuronal network activity with a time resolution of 6 minutes, and 10 seconds of human iPSC neurons exposed either to human CSF or to BrainPhys™. (**B**) Diagrams showing the proportion of experiments where network burst rate was 0 (= asynchronous), under five per minute, or over five per minute, recorded from human iPSC neurons exposed either to hCSF or to BrainPhys™. (**C-F**) Diagrams illustrating the change of neuronal network parameters under each condition respectively. Individual mean values and *p* values from Wilcoxon tests are shown. Experiments were conducted between 48 to 77 days *in vitro*. Mean, SD and *n* are presented in Supplementary Table 2. Note: each data point in the hCSF data set represents the activity of an individual human iPSC-neuronal culture exposed to a hCSF sample obtained from a human individual.

While 45% of the 22 human iPSC neurons exposed to hCSF exhibited asynchronous neuronal network activity, none of the neurons exposed to BrainPhys™ displayed asynchronous activity (Figure 4B). Instead, all human iPSC neuronal cultures in BrainPhys™ exhibited highly synchronous neuronal network activity (Figure 4B). Although the network burst duration did not differ between these conditions, BrainPhys™ elicited significantly more network bursts, a higher degree of synchrony (measured as the percentage of spikes organized into network bursts), and a significant increase in spike rate (Figure 4C—F).

In summary, human iPSC neurons exposed to BrainPhys™ show substantially higher neuronal network activity compared to when they are exposed to the physiological milieu provided by human CSF.

### Separating physiological and non-physiological network burst patterns

Based on our findings, we sought to identify neuronal network parameters that distinguish physiological from non-physiological neuronal activity across experimental conditions. Plotting the mean spike rate (spikes/minute) against the mean network burst rate (network bursts/minute) for networks exposed to ion-matched aCSF, high K^+^-aCSF, BrainPhys™, and human CSF (Figure 5), revealed that neuronal cultures exposed to physiological conditions (hCSF and aCSF with hCSF-matched ion concentration) predominantly exhibited fewer than five network bursts per minute and had a mean spike firing rate below 10,000 spikes per minute. Although this approach provides a useful starting point for distinguishing physiological from pathophysiological activity in our human neuronal networks in vitro, it cannot be applied universally to MEA experiments, as the mean spike firing rate strongly depends on the number of active electrodes used.

**Figure 5.**
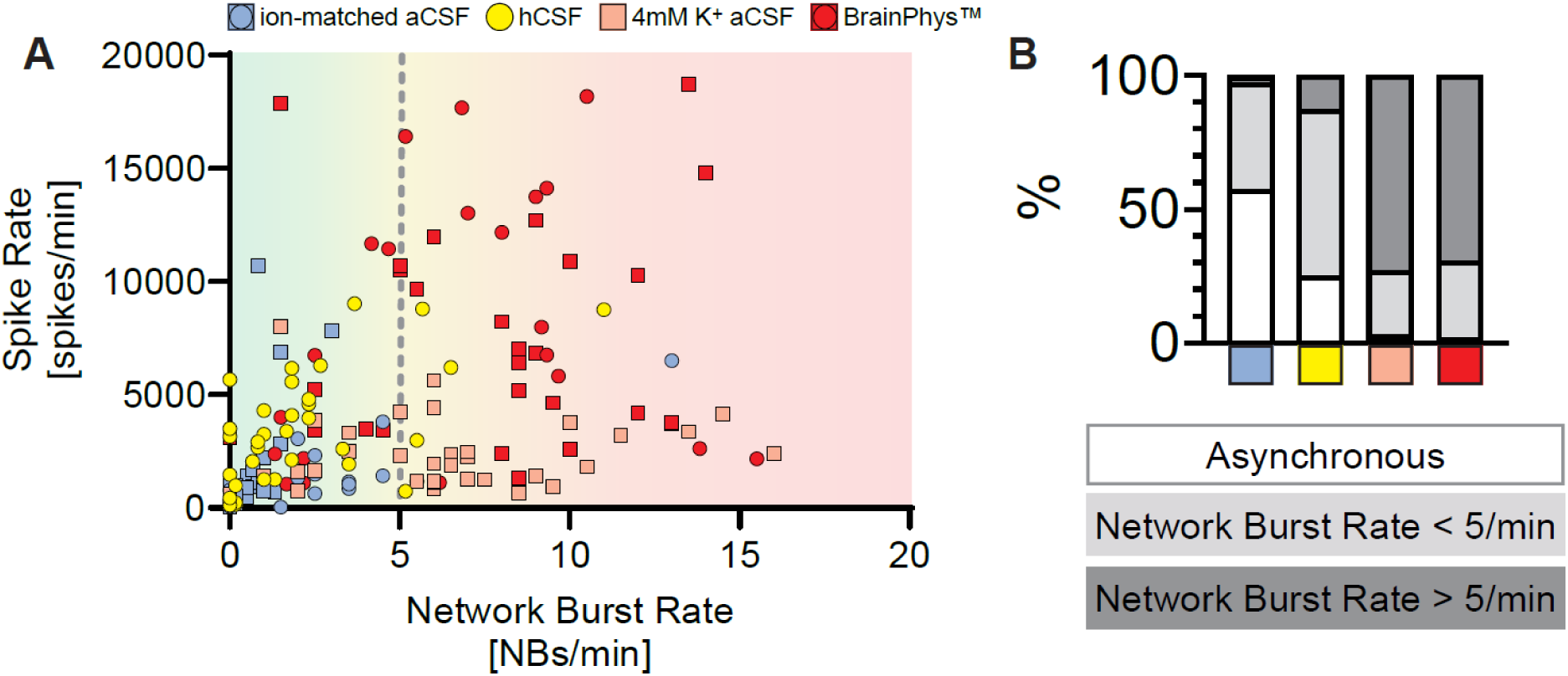
Human neuronal network activity under physiological and non-physiological ion concentrations shows specific neuronal population burst characteristics. (**A**) Mean spike rate plotted against mean network burst rate obtained from individual human iPSC-neuronal cultures exposed either to ion-matched aCSF, hCSF samples, aCSF with 4 mM K^+^, or BrainPhys™. (**B**) Diagrams showing the proportion of experiments where network burst rate was 0 per minute (= asynchronous), under five per minute, or over five per minute, under each conditions respectively.

## Discussion

Starting with the first successful *in vitro* culture of neurons by Ross Granville Harrison in 1907, through to the introduction of B27-supplemented Neurobasal™ by Gregory J. Brewer in 1993—which enabled long-term culture of rodent neurons without the need for astrocyte co-culture or serum—and later the development of BrainPhys™ by Cedric Bardy in 2015, which improved neuronal activity in both rodent brain tissue and human iPSC-derived neurons, neuroscience has progressively refined the composition of cell culture media to better mimic the neuronal extracellular milieu, aiming to improve the study of the principles of brain function in neuronal populations outside the brain.

However, it has yet to be thoroughly assessed whether commonly used cell culture media truly replicate a fundamental aspect of the *in vivo* neuronal environment: the ion concentrations to which neurons are exposed.

In this study we demonstrate that all currently used cell culture media for brain cell *in vitro* cultures contain non-physiological high potassium ion concentrations, and we present that non-physiological high potassium concentration is broadly used for acute electrophysiological experiments in neuroscience. In addition, elevated potassium concentrations—typically in the tens of millimolar range—are used to study depolarization-dependent processes underlying neuronal survival ^28^. As noted by others, there is enormous variability in these treatments, with durations ranging from seconds to days and concentrations spanning 3 mM to 150 mM KCl^28^. Because most studies employ 25–55 mM KCl^28^, the *in vivo* physiological relevance of using very high potassium concentrations to investigate depolarization-mediated neuronal survival has been questioned^28^. Experimentally, we show that “*just a little bit more*”—an increase of 1.1 mM potassium—profoundly affects neuronal network activity of human neurons and most likely that of neurons from other species as well. Using the Goldman–Hodgkin–Katz equation, the small change in equilibrium potential induced by raising extracellular K⁺ from 2.9 mM to 4.0 mM shifts the potassium reversal potential from –76 mV (at 2.9 mM) to –79 mV (at 4.0 mM). Even this slight depolarization can nonetheless have a pronounced effect on excitability through several mechanisms: (i) a few millivolts of depolarization (e.g., –76 → –79 mV) brings the resting potential nearer to the action-potential threshold (often around –50 to –45 mV), reducing the additional depolarisation needed to open voltage-gated Na⁺ channels so that smaller synaptic inputs can trigger spikes; (ii) the voltage-dependence of voltage-gated Na⁺ channel (Nav) activation and inactivation is very steep near subthreshold potentials, so even a 3 mV shift can move a larger fraction of Nav channels into the “window” range where they open more readily, increasing the probability of action-potential initiation; (iii) between –60 and –40 mV, a subset of Nav channels produces a small, non-inactivating “persistent” inward current (IₙₐP), and depolarizing from –76 to –79 mV slightly increases this persistent current, which can produce a slow ramp depolarization that further elevates excitability; (iv) when Eₖ becomes less negative, the driving force (Vₘ – Eₖ) during an action potential is smaller, so repolarizing K⁺ currents (through Kv and KCa channels) are reduced—this broadens each spike and diminishes the afterhyperpolarization (AHP), and because a smaller AHP shortens the refractory period, neurons can fire at higher frequencies; (v) some inward-rectifier K⁺ channels (Kir) begin to reduce their outward current as the membrane depolarizes toward Eₖ, transiently raising input resistance (Rₙ), so any given synaptic current produces a larger voltage deflection (ΔV = I × Rₙ) and amplifies small inputs; and (vi) in a neuronal network, a small depolarization in many cells can synchronize firing or recruit additional cells via recurrent excitation, meaning that a few millivolts of depolarization in the population can cascade into a large change in overall activity.

Here we show that a 1.1 mM increase in extracellular K⁺ not only boosts activity but also alters neuronal network dynamics to produce an apparent epileptiform phenotype. Consequently, *in vitro* electrophysiology studies—including those using human iPSC-derived brain cells, primary neuronal cultures, and acute brain slices from animals or humans—may yield misleading conclusions about normal micro- and mesoscale neuronal function when neurons are exposed to a non-physiological environment that induces pathological rather than physiological firing patterns. Given that we demonstrate that all major currently used cell culture media for culturing neurons *in vitro* contain potassium concentrations that elicit epileptiform activity, it implies that all *in vitro* neuronal models exhibit a predominantly epileptogenic baseline neuronal activity rather than a physiological one.

The implications of this can be broad and impactful, considering that the study of neuronal functional principles is widely conducted in in vitro experiments. This does not automatically invalidate such studies; however, it raises the concern that conclusions about the principles of brain cell function may have been drawn from neurons with aberrant electrophysiological properties. There are many possible reasons for the multitude of ion concentrations used for electrophysiological recordings, including lab tradition, vague literature on reference concentrations, and the assumption that CSF concentrations are equal to those in blood (e.g. Bardy *et al.* used an aCSF with a potassium concentration of 4.2mM^29^).

Other factors besides potassium concentration likely contribute to neuronal network activity. For instance, BrainPhys™, Neurobasal™, and DMEM/F12 contain lower concentrations of magnesium than human CSF. Magnesium is decreasing neuronal excitability^30, 31^ and acts as a blocker of the NMDA receptor^32, 33^. Thus, reducing magnesium concentration may further promote bursting hyperactivity of neurons exposed to BrainPhys™ with 0.91 mM Mg^2+^ (as compared to 1.1 mM in hCSF).

In our previous work, we demonstrated that acute application of hCSF increases neuronal activity compared to aCSF with matched ion concentrations also in acute rat hippocampal^34, 35^ and cortical slice preparations^36^, primary rodent neuronal cultures^37^, and in mouse ESC-derived neuronal networks^37^. Additionally, Wickham et al. showed that human CSF elicits higher activity in human organotypic brain slice cultures compared to aCSF with matched ion concentrations^38^. Here, we further support the role of neuromodulators in hCSF for physiological neuronal network activity, now for the first time using human iPSC-derived neuronal networks.

Complementing our previous discovery, in which we presented evidence for a brain mechanism that actively maintains CSF potassium concentration at approximately 2.9 mM—lower than the 4.1 mM found in serum^39^—our *in vitro* data presented here provide a potential explanation for *why* such a mechanism exists: to maintain neuronal activity at a more favourable level (a near- critical point^40^) by preventing runaway excitation.

### Limitations of the Study

We have used a set value of the ion concentrations although we are aware that they seem to fluctuate with the sleep-wake^41, 42^, and circadian^22^, cycles - suggesting that physiological ion concentrations in CSF may be dynamically regulated rather than to a single set value. This concept is an area of our ongoing research.

## Supplementary figures and tables

**Suppl. figure 1.**
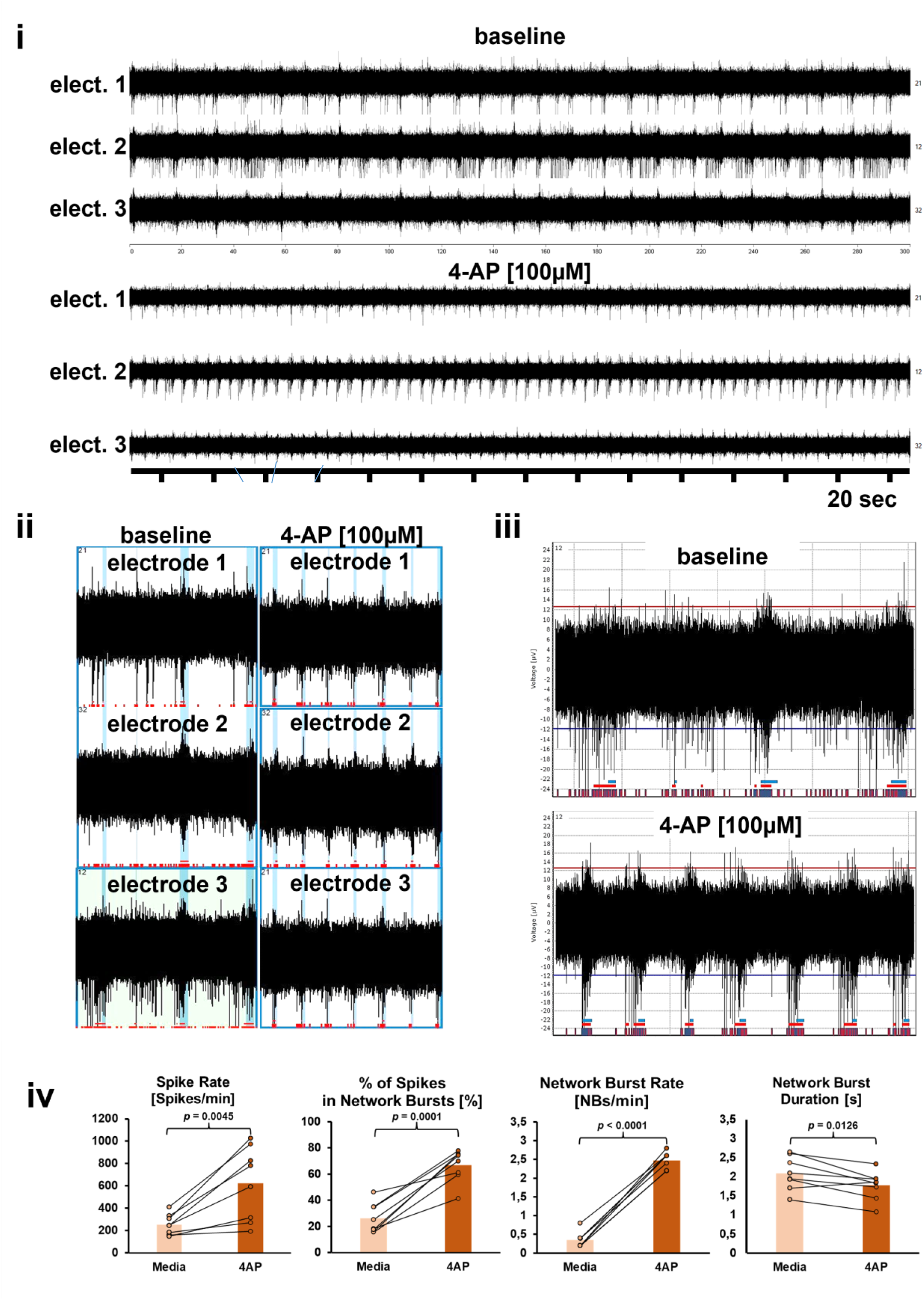
Human iPSC-derived neuronal cultures treated with convulsive compound 4-AP [100µM] show epileptiform activity. **(i)** Representative 5 minute MEA recording shows neuronal network activity before and after the application of 100 µM 4-AP detected by three electrodes. **(ii)** Representative 30 second MEA recording shows neuronal network activity before and after the application of 100 µM 4-AP detected by three electrodes. **(iii)** Depicted electrode 3 shows individual bursts detected before and after the application of 100 µM 4-AP. Note, identical 4-AP elicited epileptiform activity was confirmed in seven additional independent human iPSC-neuronal cultures (data not shown). (iv) Diagrams illustrating the change of neuronal network parameters under each condition respectively. Individual mean values and *p* values from paired *t*-tests are shown. Experiments were conducted between at 63 days *in vitro*. Note, indicated spike rates are lower as shown in figure 1-4 because the 4-AP experiments were conducted using 3D-neural aggregates cultured in 96-well MEA-plates where each well contains 3 electrodes.

**Supplementary table 1.** Ion concentrations (in mM) of applied artificial CSF obtained from peer-reviewed research articles where *in vitro* electrophysiology experiments were used. Ion concentration differences of ±15% from the measured values in human CSF assessed in this study are highlighted in red and bold.

| Na<sup>+</sup> | Cl<sup>-</sup> | K<sup>+</sup> | Ca<sup>2+</sup> | Mg<sup>2+</sup> | Used *in vitro* model for electrophysiological assessment | Reference |
| --- | --- | --- | --- | --- | --- | --- |
| 151.5 | 127.4 | 4.2 | 1.1 | 1 | Whole cell and MEA, primary rat hippocampal cultures and acute mouse brain slice | Bardy et al. PNAS; (2015) |
| 144.25 | 126.5 | 2.5 | 2 | 1 | Whole cell, acute rat brain slice | Abatis et al. Nature Neuroscience; 2024 |
| 151.25 | 133.5 | 2.5 | 2 | 1 | Whole cell, acute rat brain slice | Kraushar & Jonas, JN; (2000) |
| 141.7 | 122.3 | 3.3 | 2 | 1 | Whole cell, acute rat brain slice | Weisskopf et al. JN; (1999) |
| 146 | 127.5-130.5 | 3.5 | 2.5-4 | 3-4 | Whole cell, acute rat brain slice | Hardingham et al. JN; (2006) |
| 150 | 163 | 3 | 3 | 2 | 1. Whole cell, primary mouse hippocampal neurons | Yiu et al. Neuron; (2014) |
| 149.25 | 131.5 | 2.5 | 2 | 1 | 2. Whole cell, acute mouse brain slice |  |
| 146.25 | 126.5 | 2.5 | 2.5 | 1 | Whole cell, acute rat brain slice | Cho et al. Nature Neuroscience; (2012) |
| 151.25 | 133.5 | 2.5 | 2 | 1 | Whole cell, acute rat brain slice | Perin et al. PNAS; (2011) |
| 152.25 | 133.5 | 2.5 | 2 | 1 | Whole cell, acute mouse brain slice | Ko et al. Nature; (2011) |
| 153.25 | 133.5 | 2.5 | 2 | 1 | Whole cell, acute mouse brain slice | Franks et al. Neuron; (2011) |
| 152 | 134.4 | 4.25 | 3.2 | 1.3 | Whole cell, embryonal mouse brain slice | Yu et al. Nature; (2009) |
| 153.25 | 135.1 | 3.5 | 1.6 | 1.2 | MEA, human cortical slices | Ribierre et al. Nature Neuroscience; (2024) |
| 151.25 | 131 | 3 | 2 | 2 | Whole cell, human cortical slices | Calcagnotto et al. JN; (2005) |
| 150 | 133.8 | 3 | 1.6 | 1.3 | MEA, human cortical slices | Blauwblomme et al. Annals of Neurology; (2019) |
| 157.25 | 131 | 3 | 2 | 1 | Whole cell, acute mouse brain slice | Hsieh et al. Science Translational Medicine; (2020) |
| 157.25 | 141 | 3 | 2 | 2 | Whole cell, human brain slice | Cepeda et al. Journal of Neuroscience Research; (2003) |
| 153.25 | 135.7 | 2.5 | 4<br>(2) | 0.1<br>(1) | Whole cell, induced neurons | Cruz et al. Nature Neuroscience; (2024) |
| 150 | 153 | 5 | 2 | 2 | Whole cell, primary mouse hippocampal neurons | Wang et al. Neuron; (2016) |
| 146.65 | 153.5 | 2.5 | 2 | 1 | Whole cell, primary rat hippocampal neurons | Woo et al. Nature Neuroscience; (2009) |
| 150 | 150.6 | 5 | 2 | 0.8 | 1. Whole cell, primary mouse hippocampal neurons | Sando et al. Neuron; (2017) |
| 152 | 134.1 | 2.5 | 2 | 1.3 | 2. Whole cell, acute mouse brain slice |  |
| 146 | 130 | 3 | 2 | 2 | Whole cell, acute mouse brain slice | Sigler et al. Neuron; (2017) |
| 149.25 | 127.8 | 3 | 1.2 | 1.2 | Whole cell, acute mouse brain slice | Vardalaki et al. Nature; (2022) |
| 152.25 | 133.5 | 2.5 | 2 | 1 | Whole cell, acute mouse brain slice | Washburn et al. Nature Neuroscience; (2024) |
| 151.25 | 132 | 3 | 2 | 2 | Whole cell, acute mouse brain slice | Tomé et al. Nature Neuroscience; (2024) |
| 151.3 | 131.6 | 2.5 | 2 | 1.3 | Whole cell, acute mouse brain slice | Song et al. Nature Neuroscience; (2024) |
| 153.25 | 135.5 | 2.5 | **2** | 1 | Whole cell, acute mouse brain slice | Hattori et al. Nature Neuroscience; (2023) |
| 151.25 | 133.5 | 2.5 | **2** | 1 | Whole cell, acute mouse brain slice | Lischinsky et al. Nature Neuroscience (2023) |
| 152 | 133.5 | **3.5** | **2** | 1 | Whole cell, acute mouse brain slice | Radulescu et al. Nature Neuroscience; (2023) |
| 164.5 | 137.1 | 2.5 | **2.5** | 1.3 | Whole cell, acute mouse brain slice | Su et al. Nature Neuroscience; (2023) |
| 151.25 | 135.2 | 3 | 1.6 | **2** | Whole cell, acute human brain slice | Wickham et al. Scientific Reports; (2018) |
| 153.25<br>150 | 136.5<br>130 | 2.5<br>**4.25** | **2**<br>**3** | **2**<br>1 | 1. Whole cell, acute mouse brain slice<br>2. MEA, acute rat brain slice | Hájos et al. European Journal of Neuroscience; (2009) |
| 151.25 | 131.4 | 3 | 1.2 | 1 | Extracellular recording, human neocortical slices | Cunningham et al. Epilepsia; (2012) |
| 152.25 | 139.2 | 3 | 1.6 | **2** | Extracellular recording, human neocortical slices | Sandow et al. Frontiers in Neurology; (2015) |
| 150 | 136 | **4** | **2** | **2** | Extracellular recording, human brain slice | Huberfeld et al. Nature Neuroscience; (2011) |

**Supplementary table 2.**
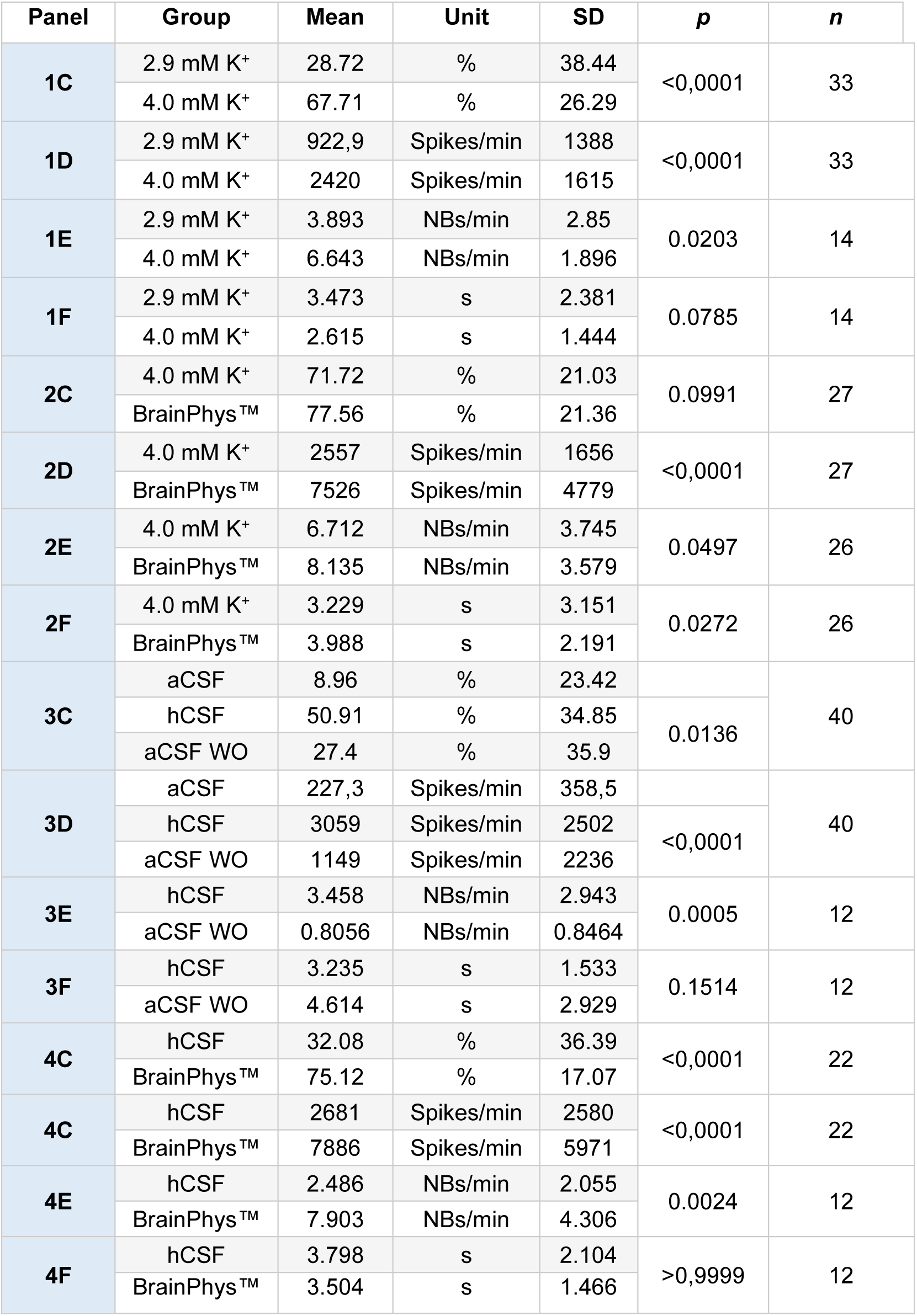
Mean-, standard deviation (SD)-, *p-*values and number of technical replicates (*n*) for Figures 1-4.

**Supplementary table 3.** Ion concentration (in mmol/L) composition of applied human CSF in the presented study.

| **Healthy control** | **Na<sup>+</sup>** | **K<sup>+</sup>** | **Cl<sup>-</sup>** | **Ca<sup>2+</sup>** | **Mg<sup>2+</sup>** |
| --- | --- | --- | --- | --- | --- |
| **1** | 145 | 2.89 | 124 | 1.21 | 1.14 |
| **2** | 147 | 2.93 | 126 | 1.17 | 1.11 |
| **3** | 146 | 2.78 | 125 | 1.16 | 1.13 |
| **4** | 145 | 2.89 | 123 | 1.14 | 1.12 |
| **5** | 145 | 2.93 | 125 | 1.16 | 1.11 |
| **6** | 147 | 2.94 | 127 | 1.20 | 1.14 |
| **7** | 148 | 3.04 | 126 | 1.16 | 1.13 |
| **Mean** | **146** | **2.91** | **125** | **1.17** | **1.13** |

## Abbreviations

aCSF: artificial cerebrospinal fluid
hCSF: human cerebrospinal fluid
iPSC: induced pluripotent stem cell
MEA: microelectrode array

## Acknowledgment

The authors would like to acknowledge grants from Alzheimerfonden (AF-556051, AF- 744871 to SI), the Åhlén Foundation (mB16/h16 to SI), Åke Wibergs Foundation (M17-0265 to SI), the Fredrik and Ingrid Thuring Foundation (2016-00225 to SI), the Psychiatry Research Foundation in Gothenburg (SI), Magnus Bervall Foundation (2019-03512 to SI), Forska Utan Djurförsök (N2023-0007 to SI), the Swedish Research Council (VR 00986 to EH and #2023-00356, #2022-01018 and #2019-02397 to HZ), Hjärnfonden (FO2021-0048 to EH), Swedish State Support for Clinical Research (ALFGBG 427611 to EH and ALFGBG- 71320 to HZ) and the European Union’s Horizon Europe research and innovation programme under grant agreement No 101053962 (to HZ).

## Declaration of generative AI and AI-assisted technologies in the writing process

During the preparation of this work the authors used ChatGPT for proof reading of the language used in the manuscript. After using this tool, the authors reviewed and edited the content as needed and take full responsibility for the content of the published article.

## Author Contribution

Conceptualization, **S.I.**; Methodology, **T.L.**, **J.I. E.A.** and **S.I.**; Investigation, **T.L.**, **J.I., M.F, K.J.**, **M.A**. **H.Z.**, **P.W.**, and **S.I.**; Formal Analysis, **T.L.** and **S.I.**; Writing – Original Draft, **S.I.**; Writing – Review & Editing, **T.L.** and **E.H.**; Funding Acquisition, **H.Z.**, **E.H.** and **S.I.**; Resources, **E.H.** and **S.I.**; Supervision, **H.Z.**, **P.W.**, **E.H.** and **S.I.**.

